# Precision Neuromodulation with Real-Time Brain Decoding for Working Memory Enhancement

**DOI:** 10.1101/2025.06.27.662056

**Authors:** Ahsan Khan, Hongming Li, Camille Blaine, Julie Grier, Ethan Hammet, Almaris Figueroa, Sarai Garcia, Romain Duprat, Justin Reber, Joseph Deluisi, Christos Davatzikos, Theodore D. Satterthwaite, Yong Fan, Desmond J. Oathes

## Abstract

Transcranial magnetic stimulation (TMS) has transformed non-invasive brain therapies but faces challenges due to variability in outcomes, likely stemming from inter-individual differences in brain function. This study aimed to address this challenge by integrating personalized functional networks (PFNs) derived from functional magnetic resonance imaging (fMRI) with a neural network-based decoder to optimize stimulation in real time during a working memory (WM) task. After identification of individualized stimulation targets, participants completed a TMS/fMRI session, performing a WM task while receiving rTMS at randomized frequencies. Decoder outputs and behavioral data during this session guided selection of optimal and suboptimal stimulation frequencies. Participants then underwent six stimulation sessions (three optimal, three suboptimal) in a randomized crossover design, performing WM and control tasks. The optimal stimulation improved WM performance by the final session, with no improvement observed in the control task. Additionally, the decoder output predicted behavioral performance on the WM task, both during the TMS/fMRI and neuromodulation sessions. These findings show that neural network-guided closed-loop neuromodulation can improve TMS effectiveness, marking a step forward in personalized brain stimulation.

**HIGHLIGHTS:**

- Closed-loop TMS guided by real-time brain decoding enhances post-stimulation behavioral effects.
- Individualized functional connectivity networks enable targeted neuromodulation
- Optimal stimulation boosted working memory performance over sub-optimal
- Brain decoder readouts predicted behavioral performance

## INTRODUCTION

Transcranial magnetic stimulation (TMS) is a non-invasive technique for modulating brain activity and has shown promise in the treatment of various psychiatric disorders including major depressive disorder (MDD), obsessive compulsive disorder (OCD) and others (McNamara et al., 2001; Mantovani et al., 2006; Carpenter et al., 2012; McGirr et al., 2021; Lisanby, 2024). Despite its clinical success, TMS outcomes remain highly variable, with only a moderate proportion of patients responding to it. For example, only half of the patients respond to TMS in MDD (Baeken et al., 2019). This has led to a considerable interest in tailoring interventions to improve treatment outcomes. One key factor contributing to this variability is individual differences in brain structure and function (Hordacre et al., 2017; Giessing et al., 2020), that influences how brain regions respond to stimulation. Additionally, the temporal dynamics of functional brain states vary both between and within individuals (Bassett et al., 2011; Calhoun et al., 2014; Finn et al., 2017), which can influence the impact of TMS on ongoing brain activity. To address these challenges, the current study combined personalized functional networks (PFNs) to identify stimulation targets, along with a neural network-based decoder for real-time and adaptive TMS delivery based on the brain state. This personalized method aimed to optimize stimulation frequency by tailoring it to each participant’s unique brain activity patterns in real time while they engaged in a Working Memory (WM) task. The effects of identified frequencies were then tested on another independent WM task during several days of neuromodulation.

The brain functions as a network of interconnected cortical and subcortical regions collaborating within a hierarchical structure (Zhou et al., 2006; Meunier et al., 2010; Doucet et al., 2011; Bassett and Siebenhühner, 2013; Park and Friston, 2013; Okamoto, 2015; Betzel and Bassett, 2016; Baum et al., 2017; De Domenico, 2017). The effect of stimulation on a target cortical region thus propagates throughout the brain resulting in network-level modulation (Chen et al., 2013; Seguin et al., 2023). Accurately determining the cortical stimulation site is therefore crucial for inducing network level changes. Various methods have traditionally been used including 5cm rule and beam F3 method (Trapp et al., 2020), however, they do not account for individual differences in brain structure and function. In contrast, functional connectivity networks (FCNs) derived from functional magnetic resonance imaging (fMRI) offer a reproducible framework for identifying individualized stimulation targets (Fox et al., 2012). Traditionally, these networks are identified using group-averaged FNCs derived from resting-state fMRI data. However, FCNs can vary markedly in their spatial topography across brain states and individuals (Ma et al., 2014; Laumann et al., 2015; Wang et al., 2015; Gordon et al., 2017; Bijsterbosch et al., 2018; Kong et al., 2018; Salehi et al., 2018). Recent studies indicate that individualized, sparse, non-negative PFNs provide improved characterization of the functional brain at the individual level (Li et al., 2017; Keller et al., 2023). Additionally, task-based fMRI enhances both within-subject and between-subject estimates of stable brain network representations compared to using resting state fMRI only (Finn et al., 2017). Considering these factors, our study utilized both resting and task-evoked brain imaging data and PFNs to identify brain stimulation targets.

Besides TMS targeting, repetitive TMS (rTMS) frequency has clear neurobiological consequences (Allen et al., 2007). Without directly measuring brain response to rTMS in humans, there remains substantial unexplained variability and even a sizable number of participants who exhibit ‘excitatory’ reactions to putative inhibitory stimulation and vice versa (Hamada et al., 2013), suggesting that common rTMS excitatory frequencies vary in their cortical effects across individuals. Currently, no systemic solution is available to effectively tune TMS frequency for optimal brain stimulation of an individual brain. We expect that tuning TMS frequency based on fMRI real-time feedback could help to personalize and therefore optimize stimulation effects on brain and behavior. Here we utilized brain decoding models with long short-term (LSTM) recurrent neural networks (RNN) (Hochreiter and Schmidhuber, 1997; Li and Fan, 2018) to obtain real-time brain state readouts and guide stimulation frequencies accordingly. Specifically, we extracted functional profiles from WM task fMRI data based on PFNs, which served as features to build brain decoding models. LSTM RNNs were adopted to learn decoding mappings between the functional profiles and brain states. This approach builds on our previous work, where we demonstrated that brain decoding models built on PFNs using LSTM RNNs can accurately decode WM and motor tasks in real time (Li and Fan, 2018).

Overall, the study investigated if using the frequency identified by the decoder and behavioral performance as optimal or suboptimal can optimize performance changes in a WM task over several days of TMS induced neuromodulation. Optimal frequencies were defined as those at which either the decoder or behavioral performance was better compared to suboptimal frequencies. It is important to note that suboptimal frequencies are only described relative to the optimal frequencies, which is intended to produce a stronger, more positive effect. We hypothesized that optimal stimulation will have a stronger positive effect on WM task performance particularly during neuromodulation sessions as compared to suboptimal stimulation and that this effect will be specific to WM and not the control task used to assess processing speed. We further expected that the decoder output would gradually improve with training and would more reliably predict how well the participants would perform during WM tasks after neuromodulation training.

## RESULTS

The study design is shown in Figure 1A. Baseline assessments included both screening questionnaires and structural/functional MRI scans during the N-back task. Subsequently, participants underwent a Pre-neuromodulation TMS/fMRI session during which rTMS at different frequencies was interleaved with N-back blocks. During this session, stimulation frequencies were classified as optimal or suboptimal based on decoder output that reflected WM network engagement and behavioral performance for each individual. After identification of optimal and suboptimal frequencies, participants underwent a crossover design receiving three consecutive optimal and three consecutive suboptimal neuromodulation sessions (six total) in a randomized order on separate days. During each session, participants performed both a WM task and a control task following rTMS stimulation. The optimal and suboptimal neuromodulation sessions were spaced at least one week apart to allow sufficient washout time for stimulation effects. The WM task used during the neuromodulation session was a delayed matching-to-sample (DMTS) task, in which participants were instructed to retain a visual stimulus in memory over a delay period and were subsequently tested on it. This task differed from the WM task used during the TMS/fMRI session to evaluate whether the effects of stimulation generalize across different WM tasks. In addition, the control task assessed processing speed and was included to test that any changes in WM were specifically attributable to the influence of stimulation on WM processes.

**Figure 1.**
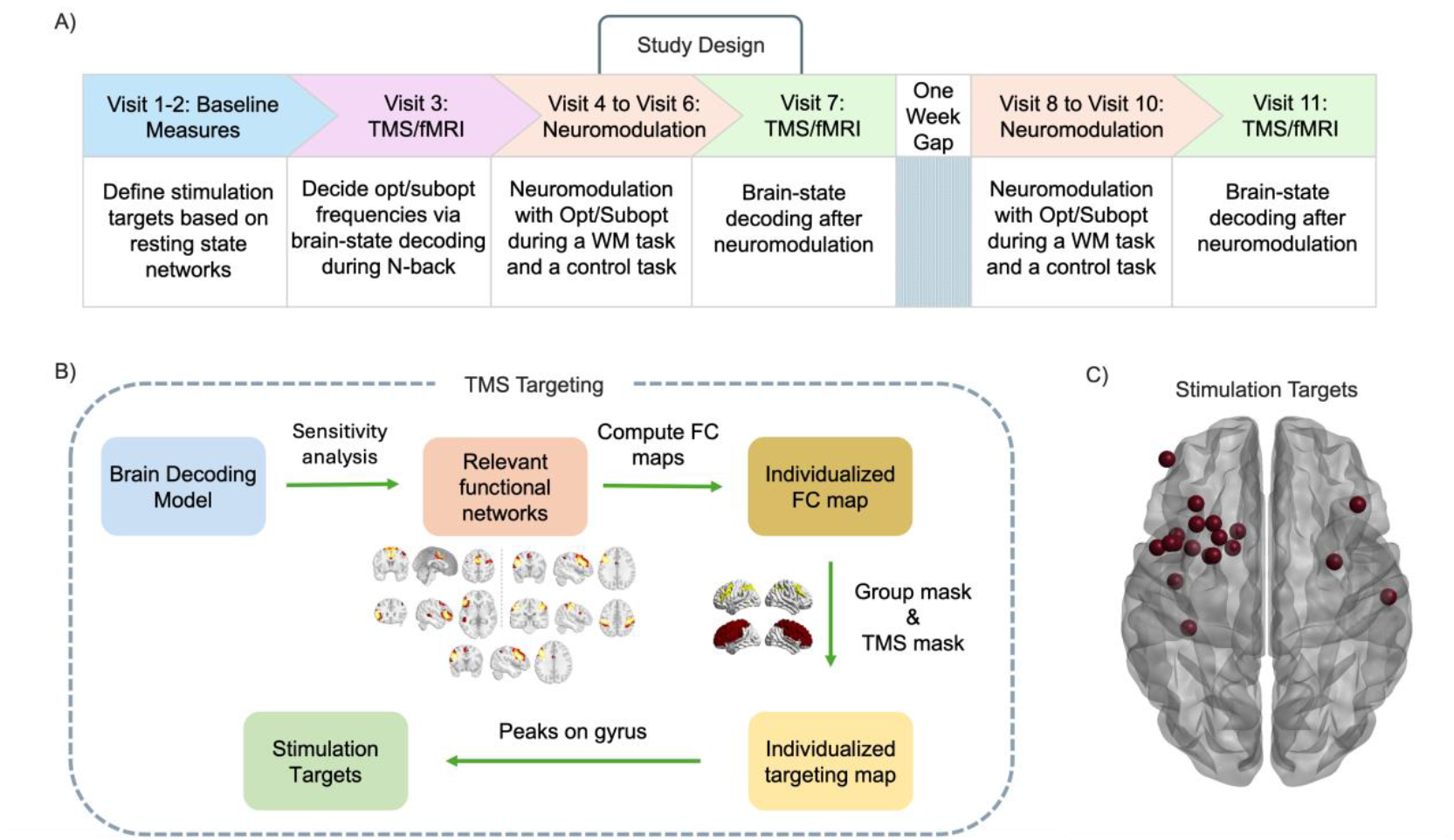
Study Design and TMS Targeting. A) Overview of the study design spanning 11 visits. B) The TMS targeting approach identifies the individualized targeting map derived from personalized functional networks (PFNs), ultimately leading to the identification of the stimulation targets for WM enhancement.(C) The final stimulation targets, individualized for each participant based on their PFN-derived maps, are shown for all individuals overlaid in MNI space.

Post-Neuromodulation TMS/fMRI sessions (Visit 7 and Visit 11) assessed the effects of stimulation and network engagement after neuromodulation training.

### Mapping Individualized Stimulation Targets

During the baseline measures, participants underwent MRI scanning to identify individualized TMS targets. We identified the stimulation target as the cortical brain region showing the strongest functional connectivity with WM-related PFNs. Specifically, we conducted a principal component analysis (PCA) on the baseline brain decoder (details provided in the methods) to determine the five PFNs most relevant to WM task decoding. These were then used to generate individualized FC maps for target identification. We then performed functional connectivity analysis on these selected PFNs to generate an individualized targeting map. The complete process for target identification is illustrated in Figure 1B, while the primary targets of all participants are shown in Figure 1C. In our study, we identified two targets for each participant to ensure that if the primary target was inaccessible for any reason with the TMS coil, the secondary target could be used as a backup (see Figure S1 for both primary and secondary targets).

### Identification of Optimal and suboptimal frequencies for WM enhancement

After identifying the individualized stimulation targets for each participant, optimal and suboptimal frequencies for stimulation were decided during a Pre-neuromodulation TMS/fMRI session. and the goal of the decoder was to predict if participant was doing a. This session consisted of two runs of the N-Back task including both a 2 back (the current item matches the one shown two trials earlier) and 0-back (the current item matches a specific target) versions. The first run, called the random loop, involved the random administration of commonly used putative excitatory rTMS frequencies (5, 10, and 20 Hz). During the random loop, the decoder predicted whether the participant was performing a 2-back task and estimated which stimulation frequency most strongly engaged the WM networks. This was reflected in the decoder output, with higher accuracy indicating greater network engagement. The second loop called the informed loop, used the information from the random loop to administer frequencies of stimulation in a specified order to test how identified frequencies engage the WM network. More details of these runs are provided in Table S1. For further analysis, we used the average decoder outputs across both runs for the optimal and suboptimal frequencies to capture overall network engagement. A schematic of the brain state decoding process is shown in Figure 2A. The N-back blocks were interleaved with three different frequencies during rTMS trains. Figure 2B illustrates the arrangement of N-back task blocks alongside the stimulation trains. Each N-back block comprised 20 trials. A limited number of pulses, only 100 in total were administered for each frequency.

**Figure 2.**
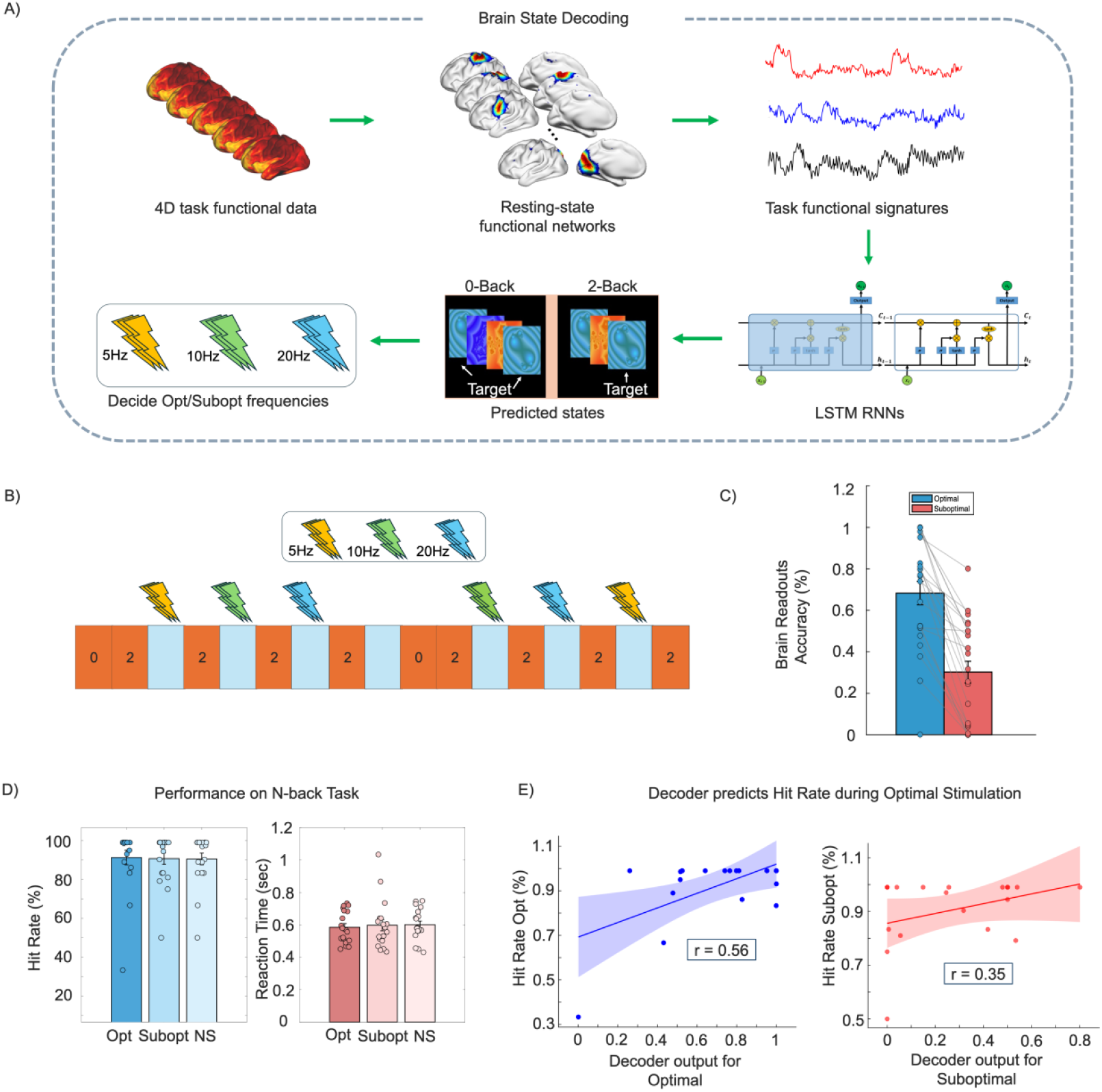
(A) Schematic of the brain state decoding approach, where the decoder distinguishes whether the participant is performing the 2-back or 0-back task. Examples of 2-back and 0-back trials used during the task are shown in the bottom panel (B) Illustration of how different stimulation frequencies were administered during the N-back task at Pre-Neuromodulation TMS/fMRI visit. Each 0 and 2 Back block represents a series of 20 trials, the decoder output and performance during N-back determined optimal and suboptimal frequencies (C) Decoder outputs, showing that the decoder predicted optimal stimulation with higher accuracy than suboptimal stimulation with individual lines (D) Average hit rate and reaction time across N-back conditions including optimal (opt), suboptimal (subopt), and No stimulation (NS). (E) The left panel shows the correlation between hit rate and decoder output during optimal stimulation, while the right panel shows the same for the suboptimal condition.

A significant difference in decoder accuracy was observed between optimal and suboptimal stimulation conditions (t (18) = 6.80, p < 0.001, Cohen’s d = 1.56), depicted in Figure 2C. However, in some cases the decoder could not reliably differentiate between the frequencies at which the accuracy fell below 50% chance. While the decoder primarily guided frequency selection, in cases where it did not clearly distinguish between optimal and suboptimal stimulation frequencies, behavioral performance on the N-back task, specifically hit rate and reaction time, was used to determine the appropriate frequency. The decision making process for frequencies as optimal or suboptimal is shown in Table S2. While the behavioral results helped guide frequency selection, the group performance on the N-back task did not significantly differ between stimulation types (optimal, suboptimal and no stimulation) for neither hit rate (F (2, 36) = 0.03, p = 0.975, η^2^ = 0.002) or reaction time (F (2, 36) = 0.71, p = 0.500, η^2^ = 0.038), as shown in Figure 2D. Surprisingly, the decoder output from the optimal stimulation condition predicted N-Back hit rate with statistical significance (r = 0.56, p = 0.013), as shown in Figure 2E. In contrast, the decoder output from the suboptimal stimulation condition did not show a significant correlation with hit rate during N-Back task (r = 0.35, p = 0.137). This finding suggests that even in the absence of a significant difference in behavioral performance, the decoder output from the pre-neuromodulation TMS/fMRI session for the optimal stimulation condition was predictive of performance, whereas the suboptimal condition was not.

### Optimal Stimulation Enhances WM Performance

Our study employed a delayed matching to sample task (DMTS) during neuromodulation sessions to assess stimulation effects on WM. In a DMTS task, participants are shown a stimulus, followed by a delay, after which they are presented with multiple stimuli and must identify the original one (a trial is depicted in Figure 3A). The delay period reflects how long participants must retain the information in WM. Common delay periods used in research, including our study, are 0 seconds, 4 seconds, and 12 seconds. The 0-second delay primarily involves discrimination and encoding processes, while longer delays (4 and 12 seconds) also engage retention mechanisms. In our study, participants underwent three consecutive optimal neuromodulation sessions and three suboptimal neuromodulation frequencies in a randomized order. Importantly, each neuromodulation session involved the administration of 2,400 pulses, compared to only 100 pulses during the TMS/fMRI session. Given this difference, we expected to observe a stronger effect on DMTS task performance during neuromodulation sessions. To assess the impact of stimulation on performance in the DMTS task, we performed a repeated measures ANOVA, with neuromodulation stimulation type (optimal vs. suboptimal), stimulation days (1 to 3), and delay (0, 4, or 12 seconds) as within-subject variables. Separate tests were conducted for reaction time and accuracy.

**Figure 3.**
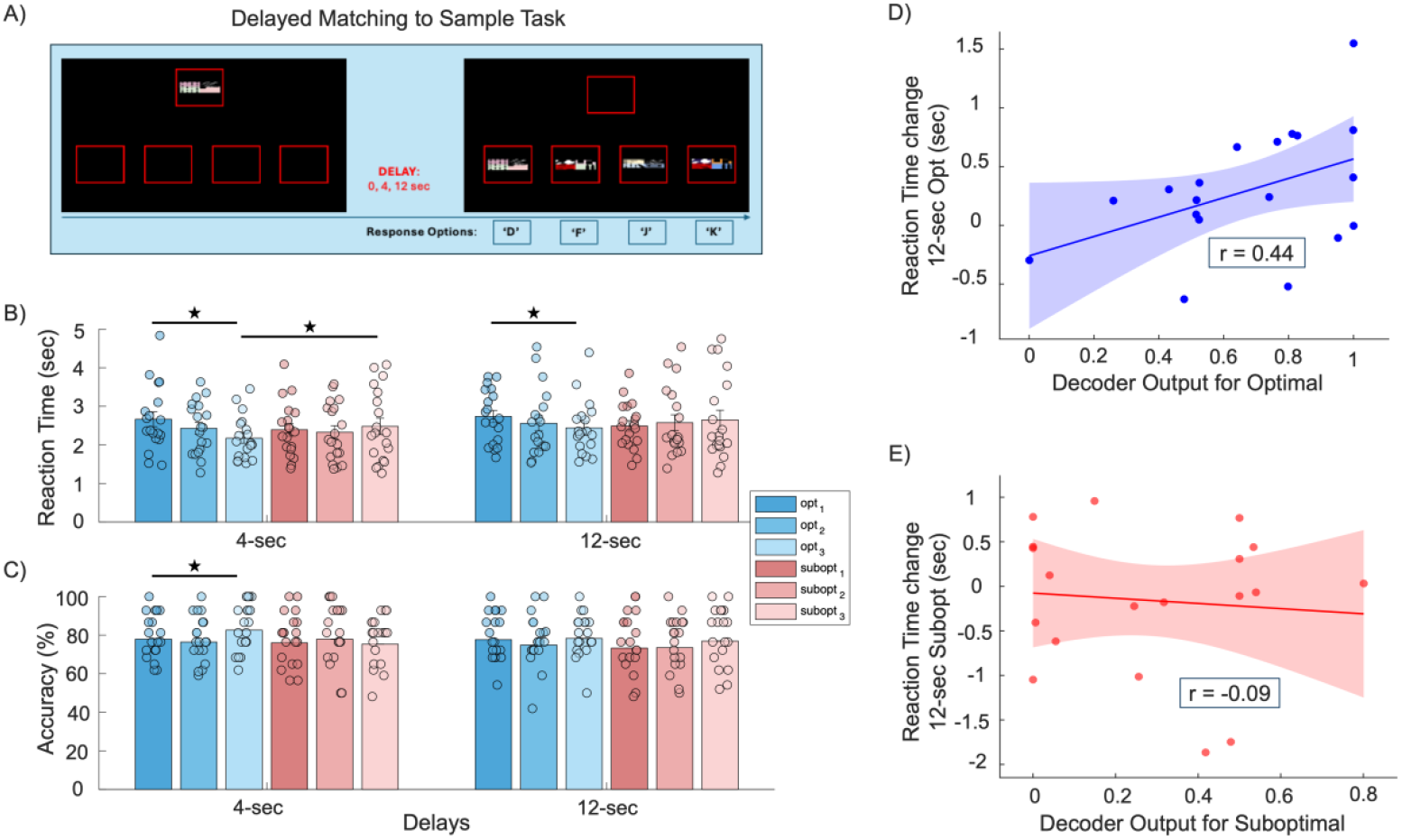
DMTS task design and performance Metrics. A) The figure illustrates the presentation of stimuli on the screen, followed by a delay of 0, 4, or 12 seconds, after which participants were required to recall the previously presented item. B) The reaction time during the DMTS task across different delay conditions is displayed, with significant differences highlighted. Participants demonstrated faster reaction times in the optimal 4-second delay condition compared to the suboptimal condition, with faster performance observed on day 3. In 12-second delay condition, participants were faster on Day 3 in optimal condition as compared to suboptimal, however no such effect was observed in suboptimal condition. The top panel presents accuracy performance across various delay periods during both optimal and suboptimal training sessions. C) The accuracy data, with a significant improvement noted in the 4-second delay condition, are shown. The blue bars represent the optimal condition, corresponding to the three days of optimal stimulation training, while the red bars represent the suboptimal condition across the three days of suboptimal training. The subscripts on “opt” and “subopt” indicate the specific day of stimulation.

For reaction time, we observed a marginally significant three-way interaction (Stimulation Type x Days x Delays) (F (4, 72) = 2.05, p = 0.097, η^2^_p_ = 0.102), a significant two-way interaction (Stimulation Type x Days) (F (2, 36) = 5.09, p = 0.011, η^2^_p_ = 0.220), and a significant effect for Delays (F (2, 36) = 30.46, p < 0.001, η^2^_p_ = 0.628). Further analysis revealed no interaction between Stimulation Type and Days for the 0-second delay condition (F (2, 36) = 0.60, p = 0.556, η^2^_p_ = 0.032), but a significant effect was found for the 4-second delay (F (2, 36) = 5.60, p = 0.008, η^2^_p_ = 0.237) and the 12-second delay (F (2, 36) = 3.33, p = 0.047, η^2^_p_ = 0.156). Post hoc analysis showed that participants were significantly faster on Day 3 compared to Day 1 during optimal neuromodulation for both the 4-second delay (t (18) = 3.31, p = 0.002, Cohen’s d = 0.758) and the 12-second delay (t (18) = 2.48, p = 0.023, Cohen’s d = 0.568). However, no such effects were observed under the suboptimal stimulation for either 4-second (t (18) = 0.55, p = 0.586, Cohen’s d = 0.126) or 12-second (t (18) = 0.855, p = 0.404, Cohen’s d = 0.196) delay condition. Additionally, the 4-second delay on Day 3 of optimal stimulation was significantly better than the 4-second delay under suboptimal stimulation on Day 3 (t (18) = 2.24, p = 0.038, Cohen’s d = 0.515). Figure 3B illustrates the significant effects observed for reaction time.

For accuracy, there was no significant three-way interaction (Stimulation Type x Days x Delays (F (4, 72) = 0.63, p = 0.644, η^2^_p_ = 0.034), nor was there a significant two-way interaction for Stimulation Type x Delays (F (2, 36) = 0.91, p = 0.413, η^2^_p_ = 0.048) or Stimulation Type x Days (F (2, 36) = 0.09, p = 0.919, η^2^_p_ = 0.005). Only main effect for Delays was significant (F (2, 36) = 11.360, p < 0.001, η^2^_p_ = 0.387). To explore whether the improvement in reaction time came at the expense of accuracy, paired t-tests were conducted on the variables that showed significant differences in reaction time. These tests indicated that participants were slightly more accurate for the 4-second delay on Day 3 of optimal stimulation compared to suboptimal stimulation (t (18) = 2.64, p = 0.017, Cohen’s d = 0.579), as shown in Figure 3C. No other comparisons reached statistical significance for either optimal or suboptimal conditions. We thus did not find evidence for a speed/accuracy trade-off in explaining the reaction time improvements specific to optimal stimulation neuromodulation. For more details on the performance metrics, including 0-delay condition see Figure S2. Overall, the results indicated that during optimal neuromodulation sessions, participants responded more quickly when recalling information after longer delays (4 and 12 seconds), which are known to engage retention mechanisms, without reduction in accuracy.

### Brain readouts from the decoder predicted performance in WM task

We further investigated whether the decoder outputs from the TMS/fMRI sessions were related to behavioral performance improvements on the DMTS task. While not statistically significant, we found that decoder output during pre-neuromodulation TMS-fMRI session showed a correlation in the positive direction with reaction time changes in the 12-delay condition from Day 1 to Day 3 (r = 0.44, p = 0.058), as shown in Figure 3D. This relationship was not observed for the suboptimal stimulation when we correlated the suboptimal decoder output with suboptimal changes in 12-second delay condition (r = −0.09, p = 0.713), as shown in Figure 3E. Future research with a larger sample size may be warranted to further explore this association.

### Optimal Frequency Stimulation Enhances Performance, Irrespective of Specific Frequency Band

Each participant received stimulation at two (optimal and suboptimal) out of three possible frequencies (5, 10, or 20 Hz) at their individualized target locations. To examine the possibility that a specific frequency was optimal across the participant cohort, we employed an exploratory Linear Mixed-Effects (LME) Model, treating Frequency as a fixed effect and Participant as a random effect to account for the unbalanced design. The three-way interaction between stimulation frequency, delay, and stimulation day on reaction time for the DMTS task was not statistically significant, (F (8, 264.57) = 1.41, p = 0.192, Cohen’s f^2^ = 0.043), suggesting that the combined influence of frequency and delay on reaction time did not differ across stimulation days. Similarly, for accuracy, the three-way interaction between frequency, delay, and visit number for the DMTS task was not significant, (F(8, 251.259) = 1.55, p = 0.142, Cohen’s f^2^ = 0.049). These findings suggest that the stimulation effects were not solely dependent on frequency regardless of whether optimization from the TMS/fMRI session was considered. The mean reaction time and accuracy with standard errors are provided in supplementary materials in Figures S3 and Figure S4 respectively. Table S2 in the supplementary materials provide information about the frequencies assigned as optimal and suboptimal for each of the participant.

### Stimulation effects were specific to the WM task

During the neuromodulation sessions, a Reaction Time Index (RTI) task was administered to assess processing speed and motor reaction time. In this task, participants were instructed to keep the ‘Reset’ button pressed and press a 2^nd^ target button when the target appeared in yellow circle (a trial is shown in Figure 4A). Reaction time was defined as the duration from target presentation to button release, while movement time referred to the duration from ‘Reset’ button release to pressing the target button.

**Figure 4.**
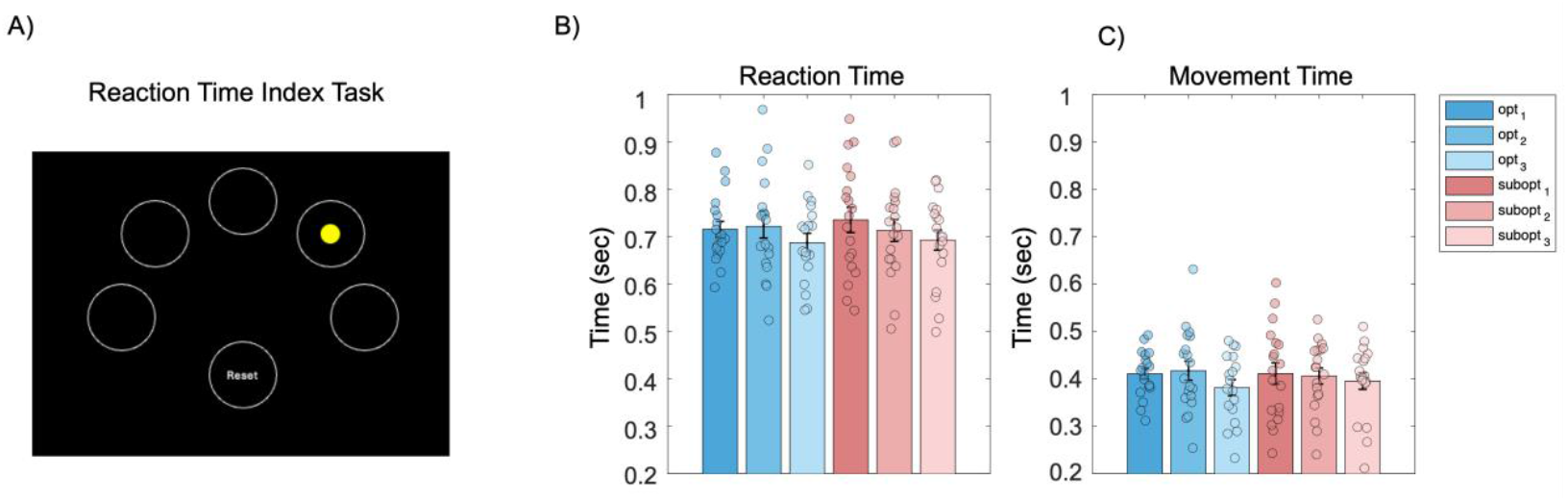
Reaction Time Index Task and performance measures. (A) This panel illustrates how the stimuli were presented on the screen. At the beginning of each trial, participants held down the reset button. When the yellow circle appeared on one of the small circles, they moved their finger to the corresponding target button. The time from the appearance of the yellow circle to releasing the reset button was recorded as the reaction time, while the time taken to move the finger from the reset button to pressing the target button was recorded as movement time. (B, C) Participants did not exhibit significant differences in performance between optimal and suboptimal stimulation conditions across any training days for either Reaction Time (B) or Movement Time (C). The blue bars represent the optimal condition, corresponding to the three days of optimal stimulation training, while the red bars represent the suboptimal condition across the three days of suboptimal training. The subscripts on “opt” and “subopt” indicate the specific day of stimulation.

The RTI task served as a control to isolate the effects of stimulation on WM, ensuring that any observed effects were specific to WM processes rather than general processing speed. A repeated measures ANOVA was conducted with neuromodulation condition (optimal vs suboptimal) and days (1 to 3) as within subject variables. No interaction between neuromodulation condition and Days were observed for neither average reaction time (F (2, 36) = 0.84, p = 0.439 η^2^_p_ = 0.045) or movement time (F (2, 36) = 0.59, p = 0.563, η^2^_p_ = 0.032). These results support the hypothesis that stimulation influenced only WM or memory retention processes, while processing speed in a non-WM related task remained unaffected. The results from the RTI task are shown in Figure 4B & C.

### Decoder brain readouts and behavioral performance after neuromodulation training

After completing the optimal and suboptimal neuromodulation sessions, TMS-fMRI sessions were conducted to assess the network engagement based on decoder accuracy and N-back task performance. Both optimal and suboptimal training session were followed by TMS-fMRI sessions involving both optimal and suboptimal frequency stimulations during N-Back task. The order of frequency administration is provided in Table S3. We expected that network engagement will be improved following neuromodulation training particularly during optimal frequency stimulation and that will reflect in improved accuracy in decoder output.

We conducted an interaction analysis using ANOVA to assess how decoder performance during the TMS/fMRI visits differed after these sessions. There was no significant interaction observed between the decoder output (optimal vs. suboptimal) and the type of neuromodulation session, F(1, 18) = 0.03, p = 0.857, η^2^_p_ = 0.002. Additionally, there was no significant main effect of stimulation, F(1, 18) = 3.70, p = 0.070, η^2^_p_ = 0.171. Nonetheless, decoder output for the optimal condition was slightly better than for the suboptimal condition, as illustrated in Figure 5A. Furthermore, there was no interaction observed in N-Back hit rate between stimulation type (optimal, suboptimal, or no stimulation) and the type of preceding neuromodulation training (optimal vs. suboptimal), F (2, 36) = 0.01, p = 0.988, η^2^_p_ < 0.001). Similarly, there was no significant interaction effect between neuromodulation training and stimulation type on reaction time, F (2, 36) = 0.31, p = 0.737, η^2^_p_ = 0.017.

**Figure 5.**
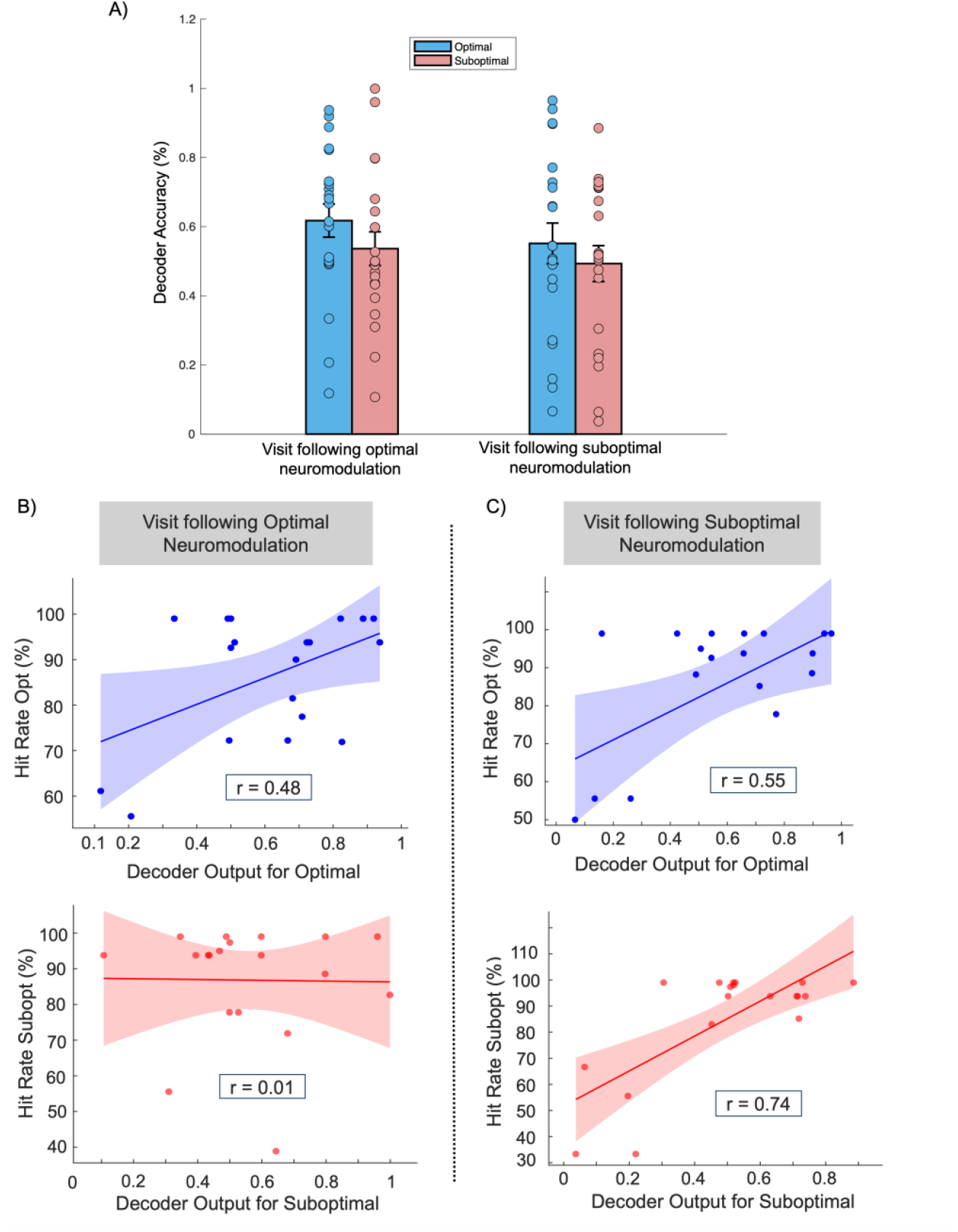
(A) Decoder output after neuromodulation: optimal stimulation is shown on the left and suboptimal on the right. The bars represent decoder activity during the TMS/fMRI session, with blue bars for optimal stimulation and red bars for suboptimal stimulation. (B) Correlation plots from the visit following optimal neuromodulation, where hit rate was significantly correlated with decoder output during the optimal TMS/fMRI session only, and not with suboptimal. (C) The visit following suboptimal neuromodulation, where hit rate correlated with decoder output during both optimal and suboptimal TMS/fMRI sessions.

We further examined the relationship between decoder output and N-back task performance. During the TMS/fMRI visit following optimal neuromodulation, N-back hit rate was significantly correlated with decoder output for optimal stimulation (r = 0.48, p = 0.037), but not for suboptimal stimulation (r = 0.01, p = 0.951), shown in Figure 5B. Interestingly, during the TMS/fMRI visit following suboptimal neuromodulation, decoder output for optimal stimulation was still significantly associated with N-back hit rate under optimal stimulation condition (r = 0.55, p = 0.016), and decoder output for suboptimal stimulation strongly correlated with hit rate under suboptimal conditions (r = 0.74, p < 0.001), shown in Figure 5C. Overall, the associations observed during the pre-neuromodulation TMS/fMRI session strengthened further in the post-neuromodulation TMS/fMRI sessions.

## DISCUSSION

This study aimed to address the variability in TMS outcomes by developing a personalized, real-time, closed-loop neuromodulation approach. By integrating PFNs derived from fMRI data with a neural network-based decoding model, we enabled adaptive, state-dependent stimulation. This approach allowed us to identify optimal and suboptimal stimulation frequencies based on decoder predictions and behavioral performance. Importantly, we found that the optimal stimulation led to significantly faster responses in a DMTS task at 4 and 12-second delay intervals, without compromising accuracy. These improvements were WM specific and stimulation did not affect control task performance, nor did frequency alone account for the effects observed in WM. Moreover, brain-behavior correlations between the decoder outputs and task performance were robust, especially during optimal neuromodulation, and improved after neuromodulation training. This suggests the decoder not only accurately tracked relevant brain states but may have refined its predictive power through repeated stimulation and training. These findings suggest that real-time brain decoding can meaningfully enhance the precision and effectiveness of TMS. By aligning stimulation with individual neural dynamics, this approach represents a promising step toward personalized and adaptive neuromodulation therapies for cognitive enhancement and clinical applications.

The primary objective of this study was to develop a real-time brain decoding framework that leverages state-of-the-art advances in PFNs derived from fMRI, combined with deep learning techniques, specifically LSTM-based RNN (Hochreiter and Schmidhuber, 1997), for adaptive brain stimulation. Utilizing functional profiles extracted from WM fMRI data based on PFNs as input features, our deep learning models learnt precise mappings between these dynamic brain states and stimulation induced brain changes (Li and Fan, 2018; Li and Fan, 2019). Importantly, we incorporated interpretable deep learning methods to rigorously identify the most informative and redundant components within our models. This approach facilitated individualized, precise targeting for TMS with concurrent fMRI, enabling real-time adaptive adjustment of stimulation parameters. Moreover, unlike most traditional network analysis techniques that rely on group-level FCN decompositions or static brain parcellations, our framework captured the dynamic variations of PFNs across both resting and task-evoked brain states (Tzourio-Mazoyer et al., 2002; Power et al., 2011; Yeo et al., 2011; Craddock et al., 2012; Zuo et al., 2014; Honnorat et al., 2015; Gordon et al., 2016). This integration of resting-state and task-based data allowed for more accurate identification of stimulation sites tailored to individual brain dynamics. Together, we developed a cutting-edge methodology for personalized and adaptive brain stimulation by integrating advanced PFN-based fMRI analysis, deep learning with LSTM RNNs, and concurrent TMS/fMRI for enhancing WM performance in healthy individuals.

During the neuromodulation training involving a WM task, where optimal versus suboptimal frequencies were tested using our approach, we found that the optimal stimulation only influenced DMTS performance in 4-second and 12-second delay trials. The DMTS captures multiple aspects of cognition, including discrimination, encoding, and retention (Paule et al., 1998). Discrimination involves recognizing the presented information, encoding refers to converting this information into an internal representation, and retention is the ability to maintain it over time. In the 0-delay condition, encoding and retention mechanisms are not engaged since there is no delay between the presentation of the to-be-remembered trial and the recall trial. However, when the delay period exceeds 0 seconds, these mechanisms come into play. The fact that we did not observe any stimulation effect on 0-second condition suggests that stimulation had no influence on the discrimination, and it only influenced mechanisms engaged in retention of WM with only 4 second and 12 second showing improved reaction time, particularly on the final day of optimal neuromodulation. Importantly, the accuracy in the optimal neuromodulation sessions was also better than the suboptimal neuromodulation session. Furthermore, the RTI task that assesses speed in the non-WM related task was not influenced by the type of stimulation, suggesting that only WM related mechanisms were modulated by stimulation. In addition, we examined whether there were frequency-specific effects by comparing performance under 5 Hz, 10 Hz, and 20 Hz stimulation during the DMTS task. The analysis did not reveal any significant effects related to stimulation frequency. While prior research has linked specific frequencies to distinct neurobiological effects, we observed that the combination of individualized targeting and frequency is what ultimately drives neuromodulatory effects. By tailoring stimulation to each participant’s unique brain region, we likely engaged diverse network dynamics, leading to target-dependent responses to frequency.

Surprisingly, no stimulation effect was evident in the N-back task during Pre-neuromodulation TMS/fMRI sessions. Although this finding is unexpected, several underlying factors could plausibly account for this, including differences in TMS dosage, and a potential ceiling effect in the N-back task. During the TMS/fMRI sessions, only 100 pulses of rTMS were administered at a specific frequency, compared to the 2400 pulses delivered in each neuromodulation session. The reduced dosage in TMS/fMRI sessions was due to time constraints, as the primary goal of those sessions was to engage the WM networks. This lower stimulation dose may have contributed to the lack of observable effects on N-back performance. Moreover, participants performed exceptionally well during the N-back task, with elevated hit rates suggesting a ceiling effect, leaving little room for measurable improvement. These factors collectively may have contributed to the absence of a detectable stimulation effect during the N-back task. Interestingly, the decoder output was significantly correlated with N-back performance at Pre-neuromodulation TMS/fMRI visit during optimal stimulation and not during suboptimal stimulation. These findings indicate that optimal stimulation possibly led to more WM network recruitment even if there were no observable changes in behavior

An interesting and unexpected finding from our study was that the decoder performance improved for suboptimal stimulation after neuromodulation training, whereas it plateaued for optimal stimulation. This effect can potentially be explained through the interplay of Hebbian and Homeostatic plasticity. Hebbian plasticity strengthens connections between neurons as they fire together, while homoeostatic plasticity prevents excessive synaptic changes (Fox and Stryker, 2017). During the pre-neuromodulation TMS/fMRI visit, the brain is responsive and the optimal stimulation possibly results in Hebbian-like plasticity by engaging WM network that reflects in higher decoder accuracy. However, once this optimized state was reached, the homeostatic mechanisms may have prevented further engagement of WM networks. In contrast, due to low plastic gain during suboptimal stimulation at Pre-Neuromodulation TMS/fMRI visit, the threshold for further plasticity may be lower, resulting in greater plastic shifts when tested again after neuromodulation training. In addition, there was no correlation between decoder output for suboptimal stimulation and N-back performance during suboptimal stimulation after optimal neuromodulation. This absence may indicate that once the brain is brought into a more efficient or selective state through optimal neuromodulation, it becomes less responsive to suboptimal stimulation frequencies, potentially reflecting a tuning of functional networks to optimal frequency. We acknowledge that these interpretations regarding what the decoder may be capturing are speculative, and we currently lack strong empirical evidence to fully support the proposed models. Nevertheless, this remains a promising direction for future investigation.

While this study introduces an innovative approach to personalizing brain stimulation using real-time neural readouts, it has several limitations that warrant consideration. First, although the decoder correlated with behavioral performance at the group level, it did not reliably predict the optimal and suboptimal stimulation frequencies at the individual level on the day of the experiment. As a result, behavioral performance remained necessary for determining stimulation efficacy. Nonetheless, the study successfully demonstrated that brain activity can be decoded in real time, a key step toward developing a fully predictive neural decoder. Second, the lack of a control neuromodulation condition limits our ability to directly assess how optimal and suboptimal training compare to a true baseline. Third, while stimulation effects were measured across multiple days, the long-term stability and durability of these changes remain unknown. Future research should assess whether the cognitive and neural benefits of optimal stimulation persist over longer periods, such as weeks or months. Finally, the study was conducted exclusively with healthy participants, so caution is needed when extending these results to clinical populations, whose baseline neural activity and responsiveness to TMS may differ significantly. Expanding the sample to include a larger and more diverse group, particularly individuals with clinical conditions, will be essential for evaluating the broader applicability and effectiveness of this approach.

In summary, we developed a precision neuromodulation approach tailored to each individual’s neural activity. By demonstrating that stimulation can be tailored dynamically based on ongoing brain state, the study provides a new avenue for optimizing cognitive enhancement and clinical interventions. In future research, this framework can potentially be extended to other cognitive domains and neuropsychiatric conditions, possibly providing more effective, precise and patient specific treatments.

## STAR★METHODS

Details method are provided in the online version of this paper and include the following.

- Key Resource Table
- Resource Availability
  - Lead Contact
  - Materials Availability
  - Data and Code Availability
- Experimental Model and Subject Details
- Methods Details
  - Cognitive Assessments
  - MRI Acquisition
  - Simultaneous TMS/fMRI Session
  - Neuromodulation Session
  - Brain Image Processing
  - Personalized Functional Networks
  - Real-Time Brain State Decoder
  - Individualize TMS Targeting
- Additional Resources

## ACKNOWLEDGMENTS

The study was supported by National Institutes of Health/Mental Health grant R01 MH120811 to DJO and YF as well as the Hart Fund in Cognitive Neuroscience (DJO). Additional support was provided by the Penn-CHOP Lifespan Brain Institute, the AE foundation, and The Penn AI2D Center.

## AUTHOR CONTRIBUTIONS

Conceptualization, DJO, YF and HL; methodology, DJO, YF, and HL; Investigation, HL, SG, CB, JD, JG, EH, AF-G; writing-original draft, AK and HL; writing-review & editing, All authors; funding acquisition, DJO and YF; resources, DJO and YF; supervision, DJO, YF, and AF-G

## SUPPLEMENTAL INFORMATION

Supplemental Information can be found online at https://doi.org/…

## STAR★METHODS

## KEY RESOURCES TABLE

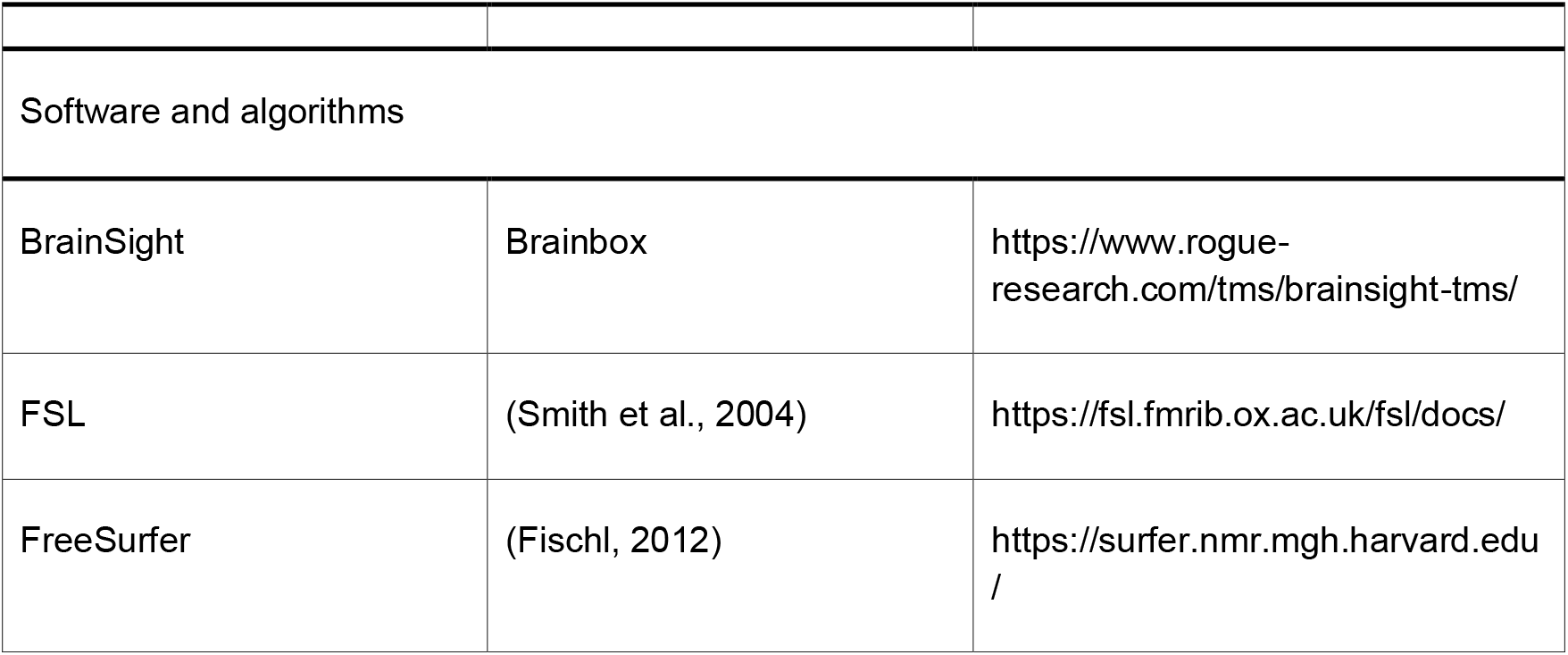
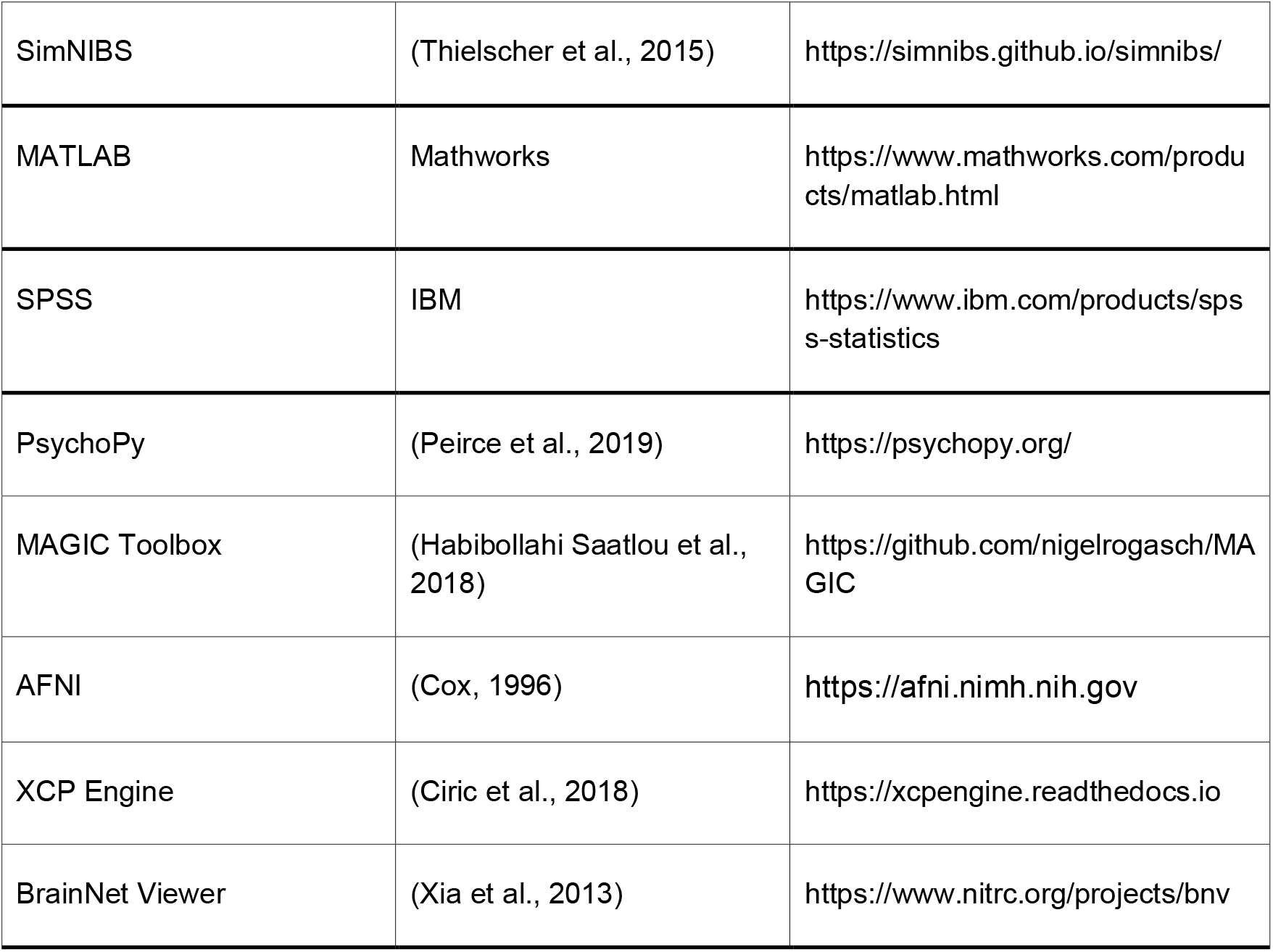

## RESOURCE AVAILABILITY

### Lead Contact

Further information and requests for resources should be directed to and will be fulfilled by the lead contact, Desmond Oathes.

### Materials Availability

Data used in this study can be requested from the lead contact, Desmond Oathes.

### Data and Code Availability

The code for computing individualized functional networks is available at https://github.com/hmlicas/Collaborative_Brain_Decomposition and https://github.com/ZaixuCui/pncSingleFuncParcel/tree/master/Step_2nd_SingleParcellation. The analysis code for individualized TMS targeting and brain state decoder is available at https://github.com/hmlicas/Loop_TMS.

## EXPERIMENTAL MODEL AND SUBJECT DETAILS

### Participants

A total of 27 participants were recruited for this study, with 23 participants completing all 11 study sessions. Four participants were excluded due to technical issues such as hardware failure during TMS/fMRI acquisition and software glitches resulting in incomplete data files in at least one of the 11 sessions. Final sample for analysis included 19 participants (N = 9 males). Participants were healthy right-handed adults (ages: ages: 18-38, mean age = 25.43 ± 5.71 years) with no present or prior psychiatric or neurological disorders. The study was approved by the University of Pennsylvania IRB. All participants gave written informed consent after receiving a complete description of the study according to the Declaration of Helsinki.

## METHOD DETAILS

### Cognitive Assessments

#### 1. N-Back Task

The WM task conducted during the TMS/fMRI was an N-Back task consisting of a 0-back condition and a 2-back condition. We adopted a fractal n-back version of WM task paradigm from (Satterthwaite et al., 2013), which was originally developed to investigate executive system maturation during adolescence. Each run comprised 10 blocks with each block containing 20 trials, during which fractal images were presented in a randomized order. Fractal stimuli were displayed for 600 ms, followed by an interstimulus interval of 2400 ms. Under the 0-back condition, participants were instructed to respond when the displayed fractal stimulus matched the pre-specified target stimulus. In 2-back condition, participants were required to indicate when the current stimulus matched the fractal image shown two trials prior. Between 2-back blocks, 400 ms gaps were inserted to allow for rTMS stimulation. Task design is shown in Figure 2D.

#### 2. DMTS task

The Delayed Matching to Sample (DMTS) task, adopted from Cambridge Neuropsychological Test Battery (CANTAB®; Cambridge Cognition, (Cognition, 2012)). During the task, a visual target pattern was presented for 3s. After target pattern disappears, four comparison patterns were displayed following a delay of 0, 4, or 12s. Reaction time and accuracy during the trials were used as outcome measures. Each delay condition included 13 trials, which were randomized across the different delay conditions. An example of a task trial is illustrated in Figure 3A.

#### 3. RTI Task

Reaction Time Index (RTI) task was used to assess motor and mental reaction speeds following neuromodulation. This task was also adopted from Cambridge Neuropsychological Test Battery (CANTAB®; Cambridge Cognition, (Cognition, 2012)). The task featured 6 circles on a black screen. Participants were instructed to select and hold the bottom circle using their mouse. A yellow dot flashed on one of the remaining 5 circles. Participants were required to move their mouse and click on the indicated target. A total of 40 trials were presented. A trial example is shown in Figure 4A.

### MRI Acquisition

MRI data were acquired using a 3 Tesla Siemens Prisma scanner. Baseline scans (T1-weighted and resting-state fMRI) were conducted with a standard 32-channel head coil (Erlangen, Germany). For interleaved TMS/fMRI scans, a custom “birdcage” head coil was used to accommodate the MRI-compatible TMS coil (RAPID quad T/R single channel; Rimpar, Germany) and a custom-built TMS coil holder. Two baseline resting-state fMRI scans were obtained with opposite phase encoding directions (A>>P and P>>A) using the following parameters: TR = 1355 ms, TE = 32 ms, FA = 68°, FOV = 216 mm, voxel size = 2.4 × 2.4 × 2.4 mm, 72 interleaved axial slices (no gap), and 640 volumes. Participants were instructed to keep their eyes open, remain still, and fixate on a central cross. High-resolution structural images were acquired using a multi-echo T1-weighted MPR sequence with the following parameters: TR = 2400 ms, TI = 1060 ms, TE = 2.24 ms, FA = 8°, voxel size = 0.8 × 0.8 × 0.8 mm, FOV = 256 mm, PAT mode = GRAPPA, and 208 slices.

### Simultaneous TMS/fMRI Session

TMS was administered within the MRI bore using an MRI-compatible Magventure MRI-B91 air-cooled coil connected to a MagPro X100 stimulator (Magventure, Farum, Denmark). Stimulation sites were marked on a Lycra swim cap on a participant’s head using neuronavigation immediately before the MRI session. The coil was positioned inside the MRI head coil, with the cable routed through the bore to generate a posterior-to-anterior induced current. Resting motor threshold (rMT) was determined in the MRI room by visually observing motor activity in the abductor pollicis brevis of the right hand. This accounted for potential influences from filters, cable length, and magnetic field interference. The rMT was defined as the minimum power required to elicit a motor reaction in 5 out of 10 stimulations. The motor threshold was adjusted according to the scalp-to-cortex distance (Stokes et al., 2007) using the following formula:

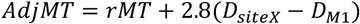

D_siteX_ is the cortical distance to the target and D_M1_ is the cortical distance to the motor cortex. Stimulation intensity for TMS was then set to 120% of adjusted MT. MRI volume acquisition (TR = 2000 ms, TE = 30 ms, FA = 75°, FOV = 192 mm, voxel size = 3 × 3 × 4 mm, 32 interleaved axial slices, 625 volumes) was synchronized with the WM task and TMS pulses using TTL triggers sent via a parallel port from a Windows PC running PsychoPy (Version 2023.1.3). A 400 ms gap was inserted between blocks of the task, allowing for two 50 pulse trains of rTMS to be delivered without contaminating the subsequent volume for each frequency. Data from runs with excessive motion (relative motion > 0.2 mm) were excluded from further analysis. A total of 600 pulses were delivered during each run of the MRI volume acquisition while participant was preforming the WM task. The order of frequencies administered were dependent on the decoder version. Results from the first TMS/fMRI session determined the frequency of TMS administered in the rest of the study sessions. The three frequencies used were 5 Hz, 10 Hz, and 20 Hz. All frequencies were administered in 50 pulse trains with a 20 second inter-train interval for 5 Hz frequency, and a 30 second inter-train interval for both the 10 Hz and 20 Hz frequency.

### Neuromodulation Session

On the participant’s first neuromodulation visit, rMT procedures were repeated in a treatment room. TMS was delivered at 110% of the AdjMT calculated using the scalp-to-cortex distance formula referred to above. Two rounds of TMS with a 5 minute break in between was administered to the participant-specific target using an MRI-Image guided stereotaxic system (Rogue Research). Each round had 24 trains of 50 pulses for the pre-determined frequency totalling 2400 pulses for a single neuromodulation visit.

Following rTMS, participants completed a RTI task and a DMTS task in a randomized order. This order was kept for all neuromodulation sessions. The tasks were practiced prior to any TMS during the first neuromodulation sessions. Pre and post-TMS behavioral surveys were administered during each visit.

### Brain Image Processing

The baseline structural MRI and functional MRI scans were pre-processed using fMRIPrep 22.1.1 (Esteban et al., 2019). Structural images underwent intensity non-uniformity correction with N4BiasFieldCorrection (Tustison et al., 2010) from ANTs 2.3.3 (Avants et al., 2008), skull-stripping with a Nipype implementation of the ANTs brain extraction workflow, and brain tissue segmentation with fast from FSL 6.0.5.1(Zhang et al., 2001). Brain surfaces were then reconstructed using FreeSurfer 7.2.0 (Dale et al., 1999). Volume-based spatial normalization to two standard spaces (MNI152NLin2009cAsym, MNI152NLin6Asym) was performed through nonlinear registration with ANTs 2.3.3.

For the functional data, a skull-stripped reference BOLD volume was generated through fMRIPrep. Head-motion parameters with respect to the BOLD reference were calculated before any spatiotemporal filtering using mcflirt (Jenkinson et al., 2002)from FSL. BOLD runs were slice-time corrected using 3dTshift from AFNI (Cox, 1996; Cox and Hyde, 1997) and resampled onto their original, native space by applying the transform to correct for head-motion. The BOLD reference was then co-registered with rigid transformations to the T1-weighted reference using bbregister (Greve and Fischl, 2009) from FreeSurfer. The BOLD time-series were resampled into standard space, generating a preprocessed BOLD run in MNI152NLin2009cAsym space. Automatic removal of motion artifacts using ICA-AROMA (Pruim et al., 2015) was performed on the preprocessed BOLD on MNI space time-series. All resamplings were performed with a single interpolation step by composing all the pertinent transformations. Post-processing procedures by Nilearn (Abraham et al., 2014), including detrend, low-pass and high-pass filtering, confounds removal (36 nuisance regressors), and spatial smoothing (6mm FWHM), were applied to the processed functional data from fMRIPrep.

### Personalized Functional Networks (PFNs)

PFNs are identified for each individual in this study to facilitate the brain readout computation and individualized target identification in the context of TMS and WM. Specifically, a spatially regularized non-negative matrix factorization technique was adopted for the identification of PFNs (Li et al., 2017), which has been demonstrated to be capable of identifying FCNs in individuals accurately while maintaining inter-individual correspondence. Since task fMRI data may provide information more sensitive to specific tasks than resting state fMRI data (Greene et al., 2018; Elliott et al., 2019; Cole et al., 2021), we concatenate resting-state and task fMRI in the temporal dimension to compute individualized FCNs. A consensus group atlas was first constructed using fMRI data of 90 healthy individuals from an inhouse dataset and PFNs were derived based on the group atlas, following the procedure in prior work (Cui et al., 2020). The number of PFNs was empirically set to 50, and one PFN was determined as a noise component by visual check and excluded from the analysis. The remaining forty-nine group-level PFNs are shown in Figure S5.

### Real-Time Brain State Decoder

To obtain fMRI based brain readout during the WM task, we developed a deep learning (DL) based brain decoder. Particularly, we utilized the LSTM RNNs to learn mappings between fMRI data and brain states, which has previously demonstrated promising performance in fMRI based decoding tasks (Li and Fan, 2019). As schematically illustrated in Figure. 2A, a predictive model of LSTM RNNs (Hochreiter and Schmidhuber, 1997) was trained based on functional signatures extracted from PFNs to classify the brain activity as 2-back state or not during the WM task fMRI session. The brain readout value was defined as the probability of being 2-back state predicted by the decoder for the time points of interest.

Given a group of *n* subjects, each having a WM task fMRI scan, and pre-computed PFNs as described previously. Based on the PFNs, the functional signatures *F*^*i*^ ∈ *R*^*T*×*K*^ used for the brain decoding were defined as weighted mean time courses of the task fMRI data within PFNs. With the functional signatures of *n* subjects, a LSTM RNNs (Hochreiter and Schmidhuber, 1997) model was built to predict the brain state of each time point based on its functional profile and temporal dependency of its preceding time points. The architecture of the LSTM RNNs consisted of two hidden LSTM layers and one fully connected layer (Li and Fan, 2019). The hidden LSTM layers were used to encode the functional information with temporal dependency for each time point, and the fully connected layer was used to learn a mapping between the learned feature representation and the brain states. The functional signatures derived from PFNs was fed into the first LSTM layer as input features, and the hidden state vector output by the first LSTM layer was used as the input to the second LSTM layer. Each LSTM layer had 128 hidden nodes. A fully connected layer with 2 output nodes was adopted for predicting the brain states as 2-back or not. Softmax cross-entropy between real and predicted brains states was used to optimize the model weights during training.

N-back WM task-fMRI data of 90 healthy participants from an independent dataset was utilized to train and evaluate the baseline decoder (90% for training and 10% for validation). Specifically, we generated training samples by splitting the functional signatures of each training participants into clip matrices of 25 by 49 with overlap of 20 time points between temporally consecutive training clips serving as data augmentation. We implemented the decoder using Tensorflow (Abadi et al., 2016). The ADAM optimizer with a learning rate of 0.001 was adopted, and the learning rate was updated every 20,000 training steps with a decay rate of 0.1. The total number of training steps was set to 100,000 with a batch size set to 32. Once the decoder was trained, it could be applied to unseen fMRI data to obtain the brain readouts for certain time points of interests.

To alleviate the effects of the discrepancy between the image characteristics of TMS/fMRI and baseline fMRI data on the decoder, we further finetune the decoder using TMS/fMRI data collected incrementally as the project proceeds, and the finetuned decoder was applied to data of incoming unseen participants. Specifically, the pre-computed individualized FCNs from baseline fMRI data for each participant was spatially registered to the native space of the TMS/fMRI data and used to extract the TMS/fMRI functional signatures. The functional signatures were then fed into the decoder to obtain the brain readouts at the time-points of interest. It is worth noting that the decoder was finetuned based on available TMS/fMRI data and only applied to incoming unseen participants to avoid data leakage.

### Individualized TMS Targeting

To identify individualized TMS target that would facilitate WM performance, we hypothesized that the potential TMS target was the brain region with highest functional connectivity to WM related PFNs. Accordingly, we first identified WM related FCNs that contributed most to the performance of WM task decoding by analysing how the changes of functional signatures of PFNs affect the baseline brain decoder. We then derived the individualized targeting map based on FC analysis of these FCNs, and finally identified the targets based on the targeting map.

Specifically, given the trained baseline WM decoder, a principal component analysis (PCA) based sensitivity analysis was carried out to identify the most significant PFNs for the WM task decoding (Koyamada et al., 2015; Li and Fan, 2019). Specifically, functional signatures of all PFNs were excluded one by one from the input and the changes in decoding accuracy on testing participants were recorded, resulting in a change matrix encapsulating changes of WM decoding performance. PCA was then applied to the change matrix to identify PCs that encoded main directions of the prediction changes with respect to changes in the functional signatures of FCNs. The top five ranked sensitive PFNs, including frontoparietal networks and dorsal attentional networks (Fig. 1B), were considered to be most relevant to WM tasks and subsequently used for target identification.

With relevant PFNs identified, the whole-brain FC map was calculated for each relevant PFN, where the FC measure for each voxel was obtained as the Pearson’s correlation coefficient between its time course and that of the PFN. The individualized FC map was calculated as the averaged FC map of all relevant PFNs. As the voxel-wise FC measures may be noisy at individual level, we restricted the candidate targets in brain regions with high FC to relevant PFNs at group level. The group-level high-FC mask was delineated by thresholding the group-average FC map of all training participants with its 90^th^ percentile. To facilitate that the target would be accessible by TMS stimulation, we further applied a TMS accessible mask during the target identification. The individualized targeting map was accordingly defined as the masked FC map by the group mask and TMS mask. The individualized target was finally identified as the peak in the targeting map located on gyrus. The gyrus region was determined by the cortical surface and sulcal depth measures generated by FreeSurfer (Fischl, 2012). Given the individualized target, E-field modeling was utilized to determine the TMS coil orientation following the procedure in (Balderston et al., 2020).

## ADDITIONAL RESOURCES

This study is registered at ClinicalTrials.gov (identifier: NCT04402294).

