## Supplementary Materials for "Precision Neuromodulation with Real-Time Brain Decoding for Working Memory Enhancement"

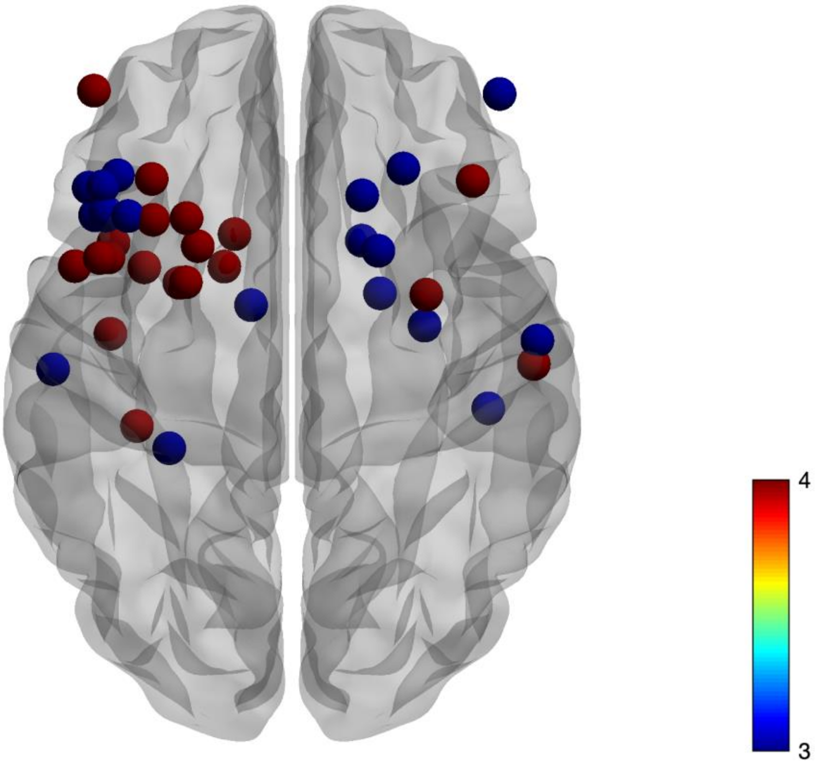


**Figure S1.** The figure illustrates the stimulation targets identified for each participant based on FCNs. Each participant has two targets: Target 1 serves as the primary target (highlighted in red), while Target 2 (highlighted in blue) acts as a backup in case the primary target is not reachable for stimulation.

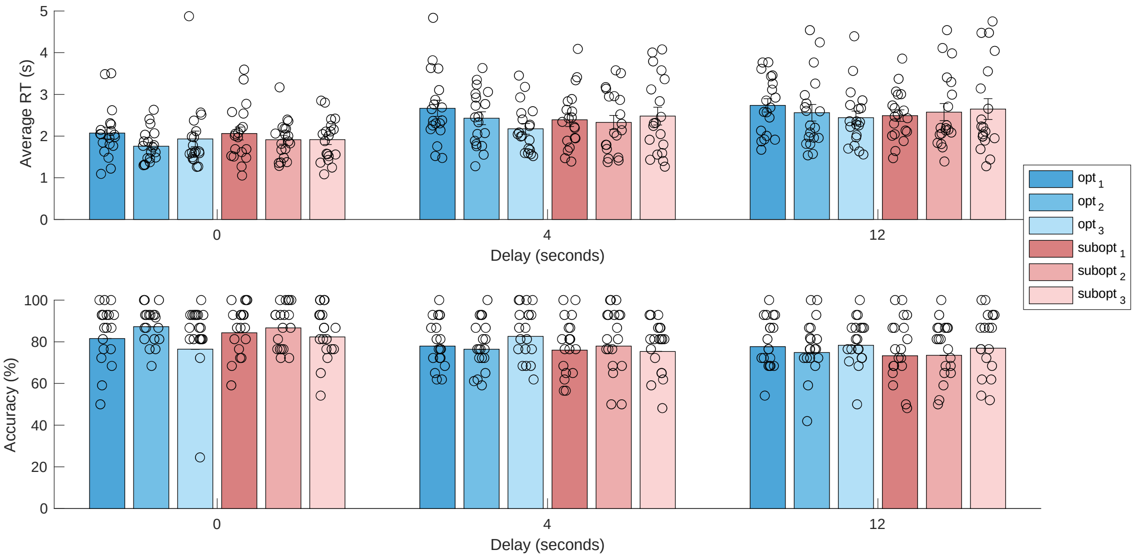


**Figure S2.** The figure shows the behavioral results for each day during DMTS task including 0-sec, 4-sec, and 12-sec delays. No significant effects were observed for 0-delay condition, while for 4-sec and 12 sec condition participants were faster on Final Day of optimal stimulation.

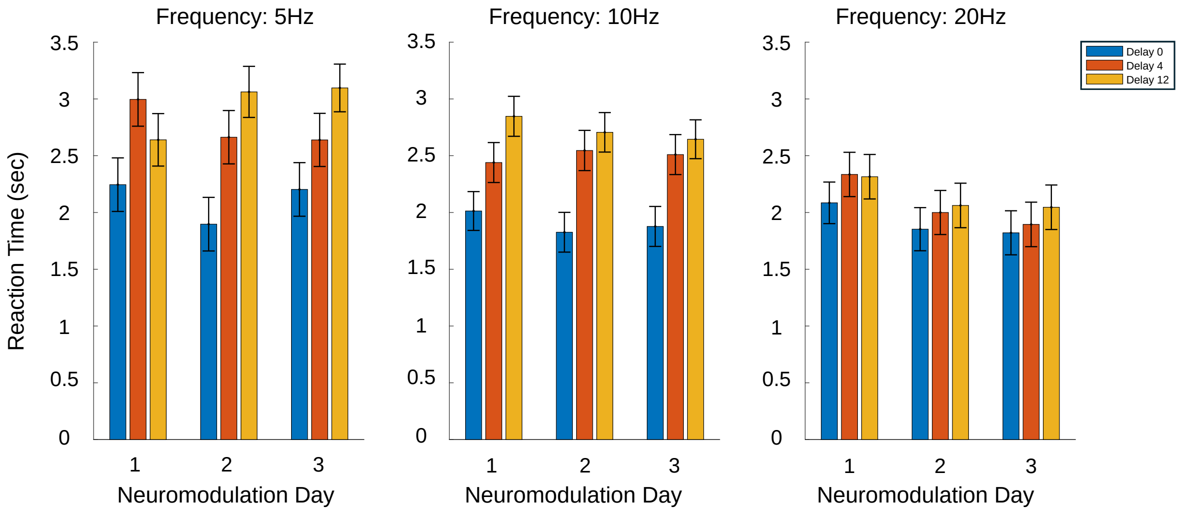


**Figure S3: Reaction Time Across Stimulation Frequencies, Delay Conditions, and Visit Numbers**

Mean reaction times (± standard error) for correct responses across three experimental visits, stratified by stimulation frequency (5 Hz, 10 Hz, 20 Hz) and memory delay conditions (0, 4, 12 seconds). Each subplot corresponds to a distinct stimulation frequency, with visit number on the x-axis and separate bars representing delay conditions. Error bars reflect the standard error of the mean. The plots illustrate interaction effects between stimulation frequency, temporal delay, and visit number on task performance, derived from a linear mixed-effects model analysis.

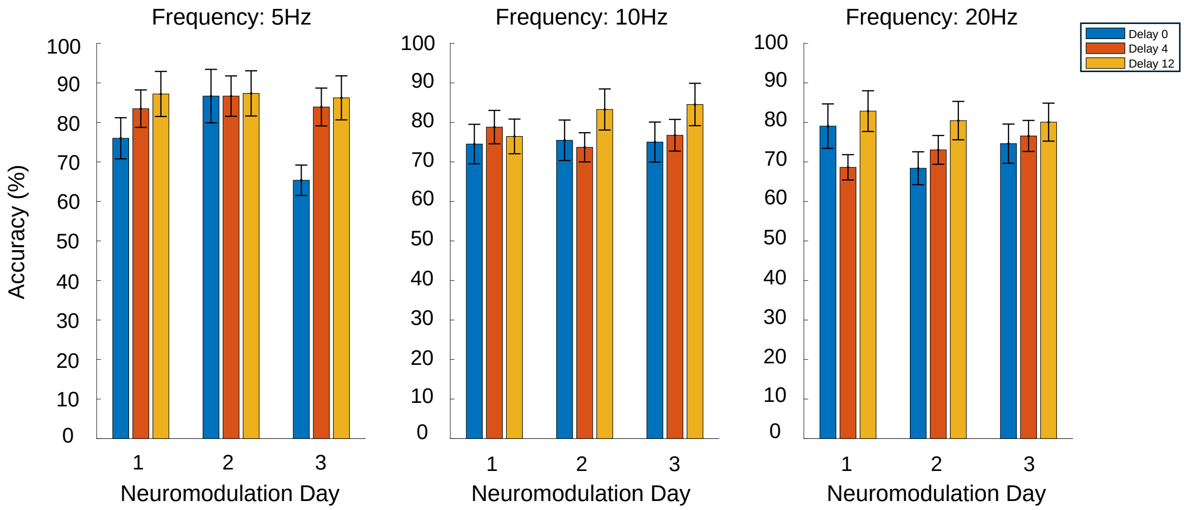


**Figure S4: Accuracy Across Stimulation Frequencies, Delay Conditions, and Visit Numbers**

This grouped bar plot illustrates mean accuracy (± standard error) across three stimulation frequencies (5 Hz, 10 Hz, 20 Hz), three delay intervals (0s, 4s, 12s), and three neuromodulation sessions. Each subplot corresponds to a frequency condition, with bars grouped by delay and color-coded accordingly. Accuracy is expressed as a percentage, reflecting performance improvement across sessions and delay conditions. Error bars represent standard errors.

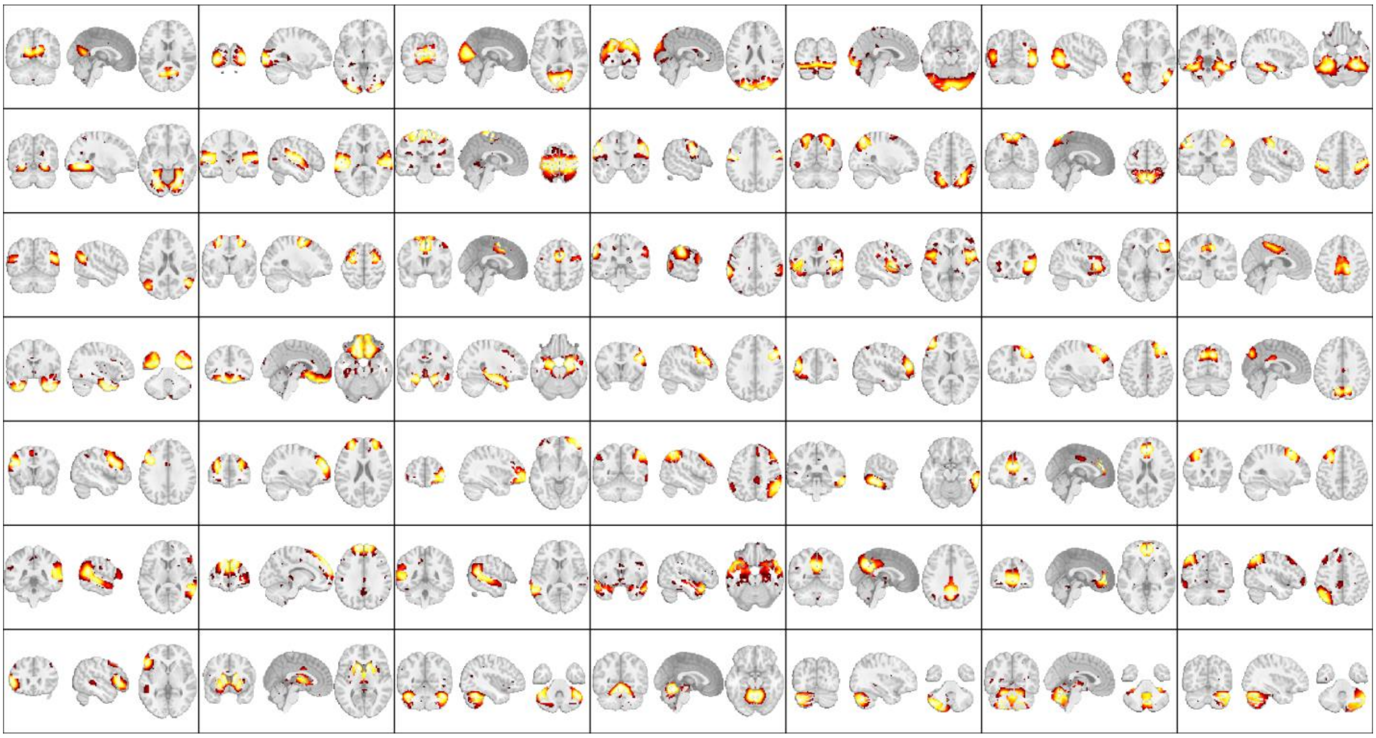

**Figure S5:** All FNs (group-level) identified by non-negative matrix factorization technique.

**Table S1**. provides the details of how the two runs of N-Back task were during Visit3.

Random Loop (random stimulation order)
no stimulation → 0-back, 2-back
first stim (**5**,10,20) → 2-back
second stim (5,**10**,20) → 2-back
third stim (5,10,**20**) → 2-back
no stimulation → 0-back, 2-back
fourth stim (5,10,**20**) → 2-back
fifth stim (5,**10**,20) → 2-back
sixth stim (**5**,10,20) → 2-back
**decides best, worst for the rest of the participant's visits**

Informed Loop (closed loop stimulation order)
no stimulation → 0-back, 2-back
first stim (**best**) → 2-back
second stim (**worst**) → 2-back
third stim (**median**) → 2-back
no stimulation → 0-back, 2-back
fourth stim (**best of 1 v. 2. v 3**) → 2-back
fifth stim (**2nd best of 1 v. 2. v 3**) → 2-back
sixth stim (**best of 4 v. 5**) → 2-back
test frequency performance

**Table S2**. provides information about the frequencies identified as optimal and suboptimal for each of the participant who completed the study.

| *Subjects* | *Optimal Frequency* | *Suboptimal Frequency* | *Visit 7* | *Visit 11* | *Decision Making* |
|---|---|---|---|---|---|
| *C106* | 10Hz | 20Hz | Optimal | Sub-optimal | Behavioral |
| *C194* | 5Hz | 20Hz | Optimal | Sub-optimal | Behavioral |
| *C236* | 20Hz | 10Hz | Sub-optimal | Optimal | Decoder |
| *C397* | 5Hz | 10Hz | Sub-optimal | Optimal | Decoder |
| *C435* | 5Hz | 20Hz | Sub-optimal | Optimal | Decoder |
| *C475* | 10Hz | 20Hz | Optimal | Sub-optimal | Decoder |
| *C527* | 20Hz | 10Hz | Optimal | Sub-optimal | Decoder |
| *C537* | 10Hz | 5Hz | Optimal | Sub-optimal | Decoder |
| *C542* | 5Hz | 10Hz | Sub-optimal | Optimal | Decoder |
| *C549* | 20Hz | 10Hz | Sub-optimal | Optimal | Behavioral |
| *C583* | 10Hz | 20Hz | Optimal | Sub-optimal | Decoder |
| *C599* | 10Hz | 20Hz | Sub-optimal | Optimal | Behavioral |
| *C605* | 10Hz | 20Hz | Sub-optimal | Optimal | Behavioral |
| *C641* | 20Hz | 10Hz | Sub-optimal | Optimal | Behavioral |
| *C694* | 10Hz | 5Hz | Sub-optimal | Optimal | Decoder |
| *C775* | 5Hz | 20Hz | Optimal | Sub-optimal | Decoder |
| *C844* | 5Hz | 10Hz | Sub-optimal | Optimal | Behavioral |
| *C905* | 20Hz | 10Hz | Optimal | Sub-optimal | Behavioral |
| *C976* | 5Hz | 10Hz | Optimal | Sub-optimal | Behavioral |

**Table S3**. provides the details of the run of N-Back task were during TMS/fMRI visit following optimal and suboptimal training.

Closed Loop (best, worst runs)
no stimulation → 0-back, 2-back
first stim (**optimal**) → 2-back
second stim (**suboptimal**) → 2-back
third stim (**optimal**) → 2-back
no stimulation → 0-back, 2-back
fourth stim (**suboptimal**) → 2-back
fifth stim (**optimal**) → 2-back
sixth stim (**suboptimal**) → 2-back
